# Application of Temperature-Responsive HIS-Tag Fluorophores to Differential Scanning Fluorimetry Screening of Small Molecule Libraries

**DOI:** 10.1101/2022.08.31.506021

**Authors:** Michael H. Ronzetti, Bolormaa Baljinnyam, Zina Itkin, Sankalp Jain, Ganesha Rai, Alexey V. Zakharov, Utpal Pal, Anton Simeonov

**Author notes:** **Correspondence:** Anton Simeonov.

## Abstract

Differential scanning fluorimetry is a rapid and economical biophysical technique used to monitor perturbations to protein structure during a thermal gradient most often by detecting protein unfolding events through an environment-sensitive fluorophore. By employing an NTA-complexed fluorophore that is sensitive to nearby structural changes in 6xHIS-tagged protein, a robust and sensitive differential scanning fluorimetry (DSF) assay is established with the specificity of an affinity tag-based system. We developed, optimized, and miniaturized this histidine-tag DSF assay (HIS-DSF) into a 1536-well high-throughput biophysical platform using the Borrelial high temperature requirement A protease (BbHtrA) as a proof of concept for the workflow. A production run of the BbHtrA HIS-DSF assay showed a tight negative control group distribution of T_m_ values with an average coefficient of variation of 0.51% and median coefficient of variation of compound T_m_ of 0.26%. The HIS-DSF platform will provide an additional assay platform for future drug discovery campaigns with applications in buffer screening and optimization, target engagement screening, and other biophysical assay efforts.

## 1 Introduction

Modern drug discovery efforts demand a toolbox with multiple assay types for the rapid high-throughput screening and confirmation of small molecule libraries to identify new therapeutic ligands. The main purpose of early-stage assays is to provide hits in an expedient and decisive path towards lead optimization. An indispensable biophysical technique for these types of screens is the thermal shift assay, also known as differential scanning fluorimetry (DSF), as it presents both a material- and cost-efficient manner of profiling engagement to a target without the need for a functional assay [1]. By utilizing a dye like SYPRO Orange that alters its fluorescence upon binding to hydrophobic patches in proteins, the unfolding dynamics of a target can be followed through a temperature gradient. Changes in the observed unfolding behavior of a target protein can be caused by ligands binding to the protein and imparting free-energy contributions that shift the Gibb’s free energy of unfolding, often seen as a stabilization of the protein in response to the temperature gradient [2, 3].

While the robustness of the DSF assay gives it a broad applicability in both sample preparation and small molecule screening, there are limitations to the assay in its current form. The environment-sensing dyes present a non-specific signal that is prone to interference from commonly employed buffer components and detergents. Additionally, the assay is optimally set up in systems with a single protein species present, whereas the presence of additional cofactors or binding partners will give rise to complex and difficult to interpret signal through the temperature ramp. In this manuscript, we describe the application of an affinity-tag fluorophore that results in a more agnostic, specific signal in a DSF setting, as well as the optimization and production of 384- and 1536-well assays to screen for small molecule binders to a model protein.

## 2 Materials and Methods

### 2.1 Protein and Reagents

*Borrelia burgdorferi* HtrA wildtype (WT) and catalytically-inactive S226A mutant (S/A) were expressed and purified as previously described [4]. The HIS-tag fluorophores RED-tris-NTA 2^nd^ Gen (NanoTemper MO-L018) and Atto-647 (Sigma 02175), were both suspended in PBS at 5 μM, aliquoted, and stored at −20 °C.

### 2.2 Casein-BODIPY cleavage assay

Protease activity was profiled using a casein substrate that has been labeled with a molar excess of BODIPY TR-X dye (EnzChek Protease Activity Kit). To construct the assay plate, 200 nL of compounds (1% DMSO v/v) and DMSO (negative control) were dry spotted into Greiner 384-well black plates (Cat #782096) and immediately mixed with 16 μL of 62.5 nM BbHtrA WT (50 nM BbHtrA WT final concentration), spun down, and incubated for 15 minutes at room temperature. Then, 4 μL of 25 μg/mL casein-BODIPY (5 μg/mL casein-BODIPY final) was added to each well, mixed, spun down, and immediately read on a Tecan Infinite M1000 in kinetic mode for 20 minutes with the following instrument settings: excitation wavelength = 590 nm ± 10; emission wavelength = 645 ± 20; gain = 95. An increase in fluorescence indicates the proteolytic liberation of fluorescent peptide fragments from the casein-BODIPY substrate, enabling comparison of proteolytic digestion rates between the DMSO, compound, and no-enzyme controls. Dose-response curves were fit using a four-parameter log logistic fit for the replicate reactions.

### 2.3 Casein digestion using SDS-PAGE

Proteolytic activity was profiled against native casein protein by SDS-PAGE to detect proteolytic fragments after incubation with BbHtrA. Briefly, 500 nM BbHtrA WT was mixed with 100 μM of small molecule (1% DMSO v/v final) for 15 minutes at room temperature. Then, casein was added to a final concentration of 25 μM and incubated for 90 minutes at room temperature. After the reaction, 15 μL of each sample was mixed with 4X LDS sample buffer (ThermoFisher #NP0007) with reducing agent (ThermoFisher #NP0009) and separated on a 10% bis-tris gel (ThermoFisher #NP0315) with MES running buffer. The gel was then washed 4 times for 5 minutes each with deionized water and stained with Imperial Blue stain (ThermoFisher #24615) for two hours before washing in deionized water overnight. The stained gel was then imaged using a ChemiDoc MP with default Coomassie Blue imaging settings.

### 2.4 Microscale thermophoresis

To evaluate the binding affinity of RED-tris-NTA and Atto-647 against HIS-tagged BbHtrA, a 16-point 1:1 serial dilution of 4 μM BbHtrA S/A was made in PBS-T (pH 7.4, 0.01% Tween-20) in a final volume of 10 μL. Then, 10 μL of a 10 nM solution of either fluorophore in PBS-T was added to each point of the BbHtrA S/A titration, mixed, and incubated in the dark at room temperature for 30 minutes. Samples were then loaded into standard capillaries and read on a NT.Automated (Nanotemper) using 10% excitation energy and “medium” MST power settings. All MST data was analyzed using MO.AffinityAnalysis software (Nanotemper) and fit using the time period +0.5 to +1.5 seconds after application of the IR laser. Data was checked for sharp capillary shapes with a single peak and consistent initial fluorescence before application of the IR laser. Normalized fluorescence values were then fit using the standard K_d_ model derived from the law of mass action with the concentration of fluorophore fixed at 5 nM.

### 2.5 HIS-tagged differential scanning fluorimetry

The Roche LightCycler 480 II and Roche LightCycler 1536 were used to run HIS-DSF assays with the 618 nm and 660 nm filters used for excitation and emission, respectively. The optimal labeling reaction was determined by creating a matrix of 7 μM 1:1 BbHtrA S/A and 500 nM 1:2 Red-tris-NTA dispensed in equal volumes onto Roche 384 well PCR plates by ECHO 550 acoustic liquid handling, incubated for 30 minutes at room temperature, and then sealed with optically-transparent seals before loading into the Roche 480 II qPCR instrument. Samples were melted in standard thermal unfolding mode with a thermal gradient from 20 to 95 °C at maximum ramping speed with 4 acquisitions per degree. The raw thermal curves were then analyzed to derive T_m_ values by the maximum first derivative method using Roche Thermal Shift analysis software.

Once optimized, a fresh mixture of the optimal BbHtrA S/A HIS-DSF labeling reaction was made for each experiment. Briefly, 3 μM HIS-tagged BBHtrA S/A was mixed with 200 nM Red-tris-NTA dye in PBS (1X, pH 7.4) and incubated for 30 minutes at room temperature. Small molecules and DMSO (negative vehicle control) were dispensed using the ECHO 550 into Roche PCR assay plates (384 or 1536-well) and spun down at 1,250 × g prior to dispensing labeled protein into the assay plate. The assay plates were sealed with clear optical foil, centrifuged again, and run immediately on a Roche 480 II or Roche 1536 qPCR instrument. Standard melting curve settings were used with a thermal gradient from 37 to 95 °C at maximum ramping speed with 4 acquisitions per degree. Data was exported from the qPCR software and analyzed using the Roche Thermal Shift analysis software.

### 2.6 Nano-Differential Scanning Fluorimetry

Real-time monitoring of the 330 nm, 350 nm, and backscattering absorbance of BbHtrA S/A samples in the presence of compound was performed using a NanoTemper Prometheus NT.48 instrument at an excitation wavelength of 280 nm. First, 10 μM BbHtrA S/A was incubated with 100 μM compound or DMSO for 15 minutes at room temperature. Then, samples were loaded into standard capillaries, loaded onto the capillary tray, and the temperature was increased from 25 to 95 °C with a ramp rate of 2.0 °C/minute. Since the small molecules interfere with the signal in the UV range of the instrument, the backscattering absorbance was used and plotted as a function of temperature. Three biological replicates were carried out for each condition, and their means and standard deviations are depicted.

### 2.7 Data analysis

All data and figures were generated using GraphPad Prism 9.

## 3 Results

### 3.1 Establishing the HIS-tag DSF Assay

To provide target engagement data in a target-agnostic manner for our screening campaign, we sought to develop a biophysical thermal-shift based method that was amenable to 1536-well high-throughput format. The assay was also intended to be adaptable to other targets and more tolerant of buffer conditions that interfere with traditional, SYPRO Orange-based thermal shift experiments. Conceivably, an affinity-tagged fluorophore that gives different quantum yields dependent on the surrounding protein structure could enable a readout on target engagement much like SYPRO-Orange DSF (Fig 1A).

**Fig 1.**
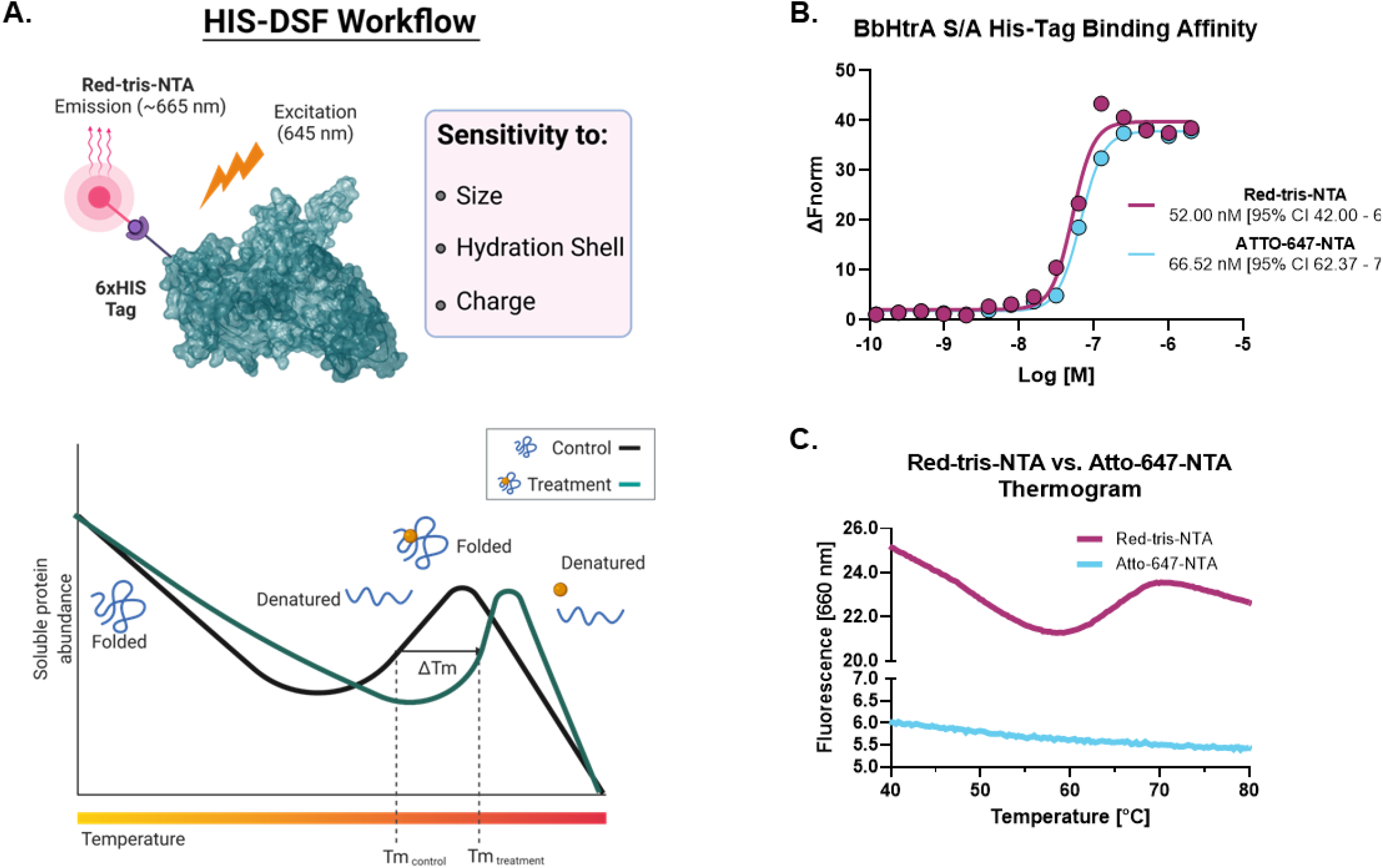
Overview of the histidine-tagged differential scanning fluorimetry assay (HIS-DSF). (A) Schematic representation of the design and workflow of the HIS-DSF assay. (B) Evaluation of the affinity for the NTA-fluorophores towards HIS-tagged BbHtrA S/A by MST. (C) Testing HIS-tag directed fluorophores in a differential scanning fluorimetry experiment. Labeled protein (as described in the *Materials and Methods* section) was run in a standard DSF melting experiment to check for the presence of melting curves.

#### Testing HIS-tag fluorophore properties and binding to BbHtrA

Toward these goals, we sought to profile a set of dyes with specificity towards HIS-tagged proteins for their performance in differential scanning fluorimetry experiments. First, the affinity of the HIS-tag fluorophores towards our target BbHtrA protein was tested using microscale thermophoresis. Both NTA fluorophores have a similar binding affinity for BbHtrA S/A, with Red-tris-NTA and ATTO-647-NTA binding to our target protein with K_d_ of 52.00 and 66.52 nM, respectively (Fig 1B). Interestingly, while Atto-647 and Red-tris-NTA demonstrate similar affinities for HIS-tagged BbHtrA S/A, only the Red-tris-NTA fluorophore can produce the typical DSF melting curve when tested in a thermal ramp (Fig 1C).

#### Miniaturization and Optimization of the DSF Assay

The ideal combination of probe and target protein for a DSF assay will give high signal-to-noise ratios and a tight distribution of melting temperatures in the negative control condition. To that end, the HIS-tag DSF assay was established by testing a matrix of protein and dye concentrations in 384-well format for the combination with the sharpest first derivative peak and cleanest thermogram. After testing triplicate 5 μL reactions of each matrix point of BbHtrA and RED-tris-NTA, the two best conditions identified were 3 μM BbHtrA and either 500 or 166.7 nM RED-tris-NTA (Fig 2A). To minimize cost of reagents, 200 nM RED-tris-NTA was chosen to move forward in continued optimization. Follow-up testing of 36 replicates of the optimized RED-tris-NTA and BbHtrA S/A reaction concentrations gave raw thermal melting curves with similar sigmoidal profiles and initial signal, that are further exemplified in the first derivative plot of these raw melting curves (Fig 2B/C). Additionally, the peaks of the first derivative of the melting curve, defined as the T_m_, are tightly grouped at 67.98 °C (95% CI 67.96 – 68.01 °C). The signal to noise ratio (S/N) for the HIS-DSF assay in 384-well format, defined as the ratio of the mean signal to the standard deviation of that signal, was calculated to be 730.8, with a percent coefficient of variation (% CV) of 0.09%.

**Fig 2.**
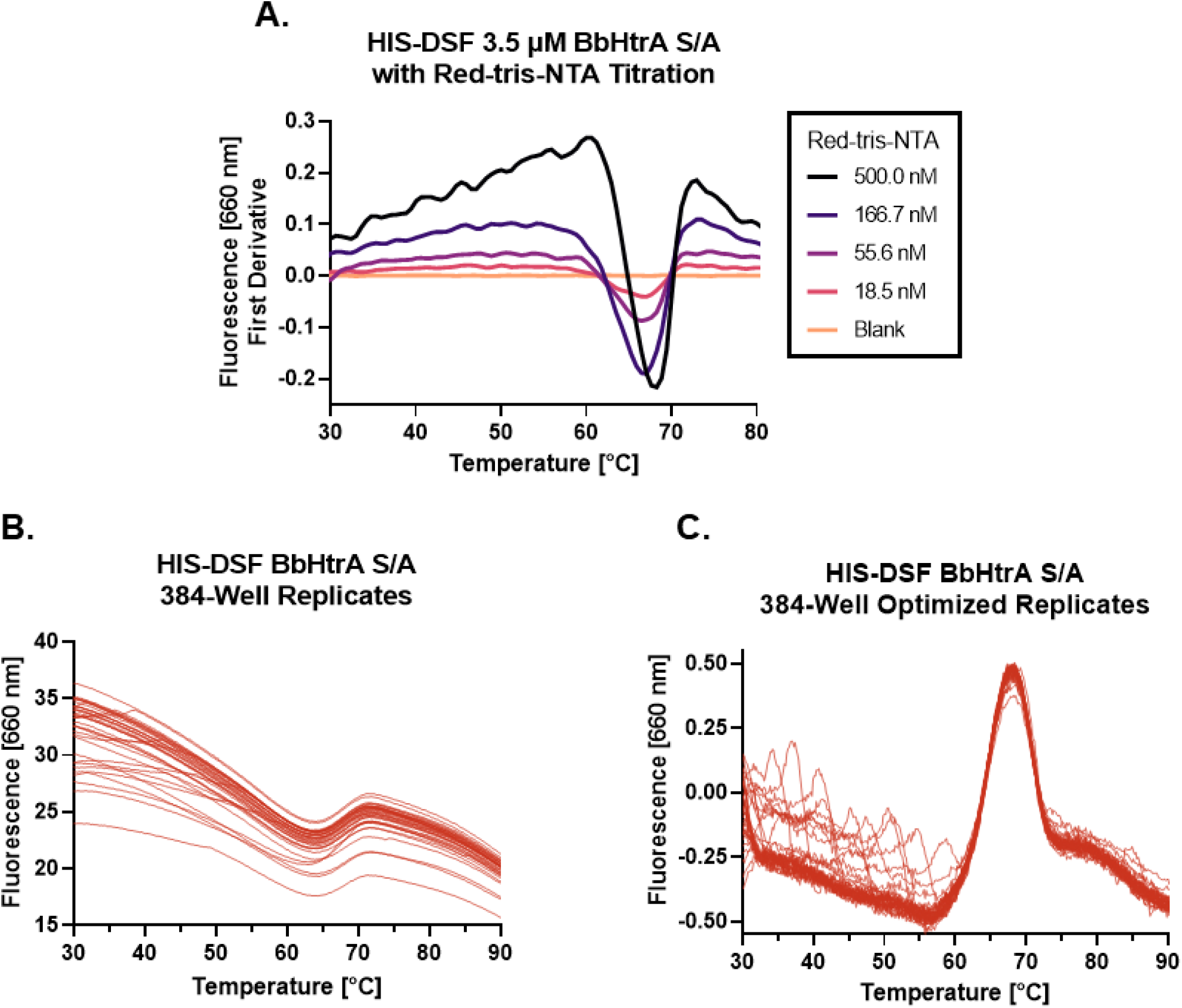
Establishing conditions for a HIS-DSF assay. (A) A matrix of BbHtrA S/A and Red-tris-NTA concentrations was profiled for its performance in a DSF temperature ramp. Pictured is a representative experiment of the 3.5 μM protein concentration group, with individual curves representing different concentrations of Red-tris-NTA or buffer. (B) Raw thermogram traces for n = 36 replicates of the matrix-derived best concentrations of protein and dye in 384-well format. (C) First derivative of the n = 36 replicate wells of the optimized HIS-DSF reaction.

We attempted to further miniaturize the assay to 1536-well format after modifying the excitation and emission filters in the Roche LightCycler 1536 for an appropriate set (excitation: 618 nm, emission: 640 nm). Miniaturization of the optimized BbHtrA HIS-DSF labeling mixture revealed that variation in T_m_ and initial relative fluorescence (RFU) signal was acceptable down to 1000 nL (Fig 3A). The effect of DMSO was profiled on the melting behavior of BbHtrA S/A with a 5% v/v 1:2 DMSO titration, simulating the solvent concentrations that would take place in a small molecule screening campaign. The thermal unfolding profile of BbHtrA S/A did not vary significantly in any of the DMSO concentrations tested, although a slight destabilization is detected from 1.67% DMSO and above (Fig 3B/C). Final testing of 36 replicates of the optimized BbHtrA HIS-DSF reaction showed an average T_m_ of 63.77 °C (95% CI 63.66 – 63.97 °C) with a S/N of 191.4 and negative control % CV of 0.52% (Fig 3D).

**Fig 3.**
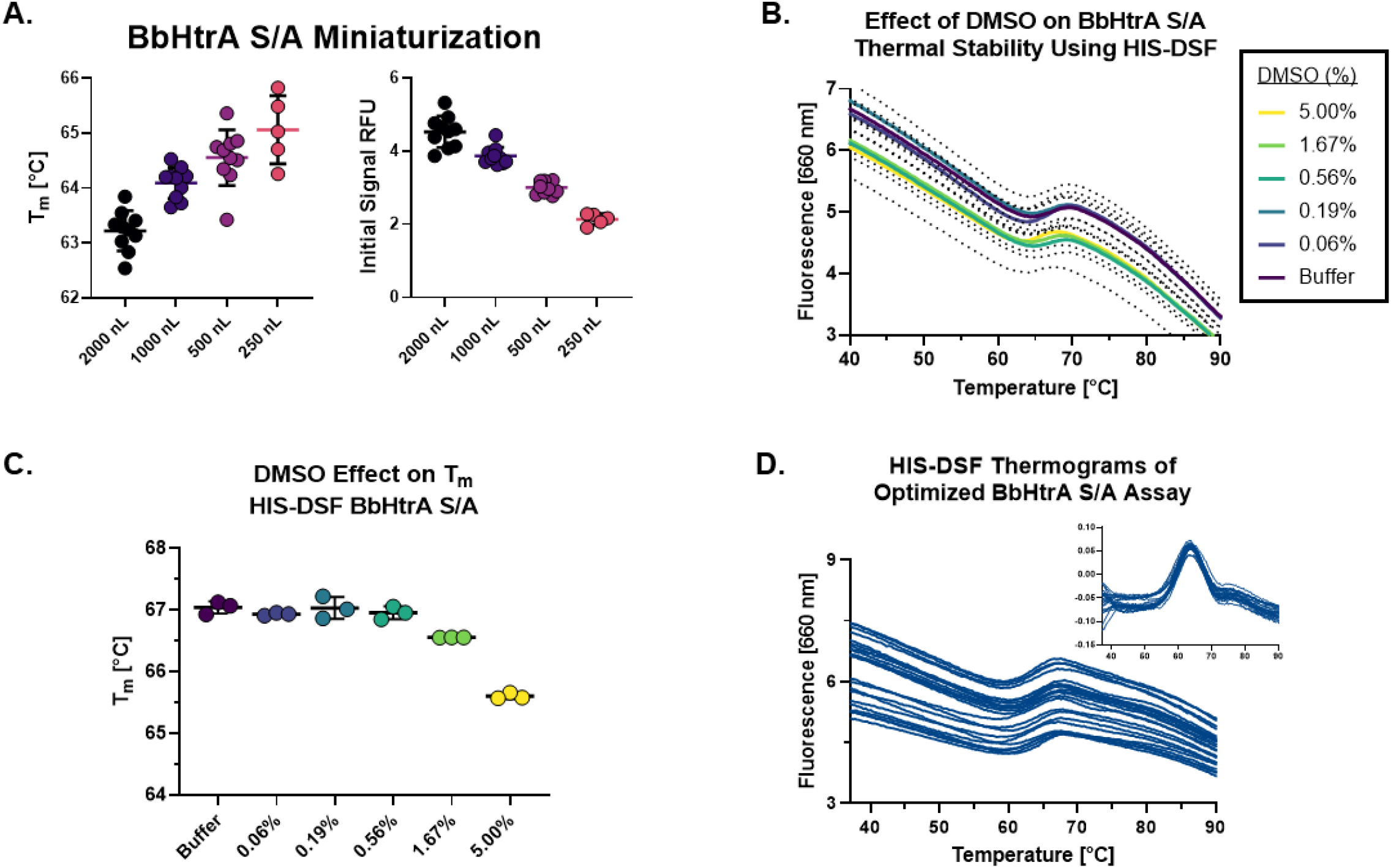
Optimization and miniaturization of the 1536-well HIS-DSF assay. (A) Scatter plot of T_m_ and initial signal RFU values from miniaturization of the HIS-DSF BbHtrA S/A reaction into 1536-well format. Lines represent the mean of three replicates with error bars representing the standard deviation. (B) Raw thermogram of the optimized HIS-DSF BbHtrA S/A reaction in the presence of different concentrations of DMSO. Lines represent the mean of three replicates with dotted lines representing the standard deviation. (C) Scatter plot of the T_m_ values from the optimized HIS-DSF BbHtrA S/A reaction in the presence of different concentrations of DMSO. Lines represent the mean of three replicates with error bars representing the standard deviation. (D) Raw thermogram traces for n = 36 replicates of the optimized and miniaturized HIS-DSF BbHtrA S/A reaction in 1536-well format. The inset graph shows the first derivatives of the raw thermograms from the same reaction.

### 3.2 Proof-of-concept screening of compound libraries and confirmation of hit molecules

#### Primary screening of a protease-targeted small molecule library

The production-readiness of the assay was tested by performing a single-dose screen in triplicate of the NCATS Protease Inhibitor library, a curated collection of 872 protease-targeted small molecules. To evaluate the robustness of our analysis on assay variation, each replicate was performed on separate days with fresh labeling reactions of BbHtrA. As there are no known ligands that thermally stabilize HtrA proteins without interfering with the HIS-tag labeling (ZnCl_2_ stabilizes HtrA proteins, inhibits the protease activity, and interferes with NTA labeling), only the negative control variation was used in selecting compounds moving forward [5, 6]. The distribution of T_m_ values for the aggregated DMSO negative control group was in line with previous values obtained during optimization, giving an average T_m_ of 63.86 °C ± 0.33 and an average % CV of 0.51%. The raw thermograms of the DMSO control group between runs also demonstrated low inter-run variability between traces (Fig 4A).

**Fig 4.**
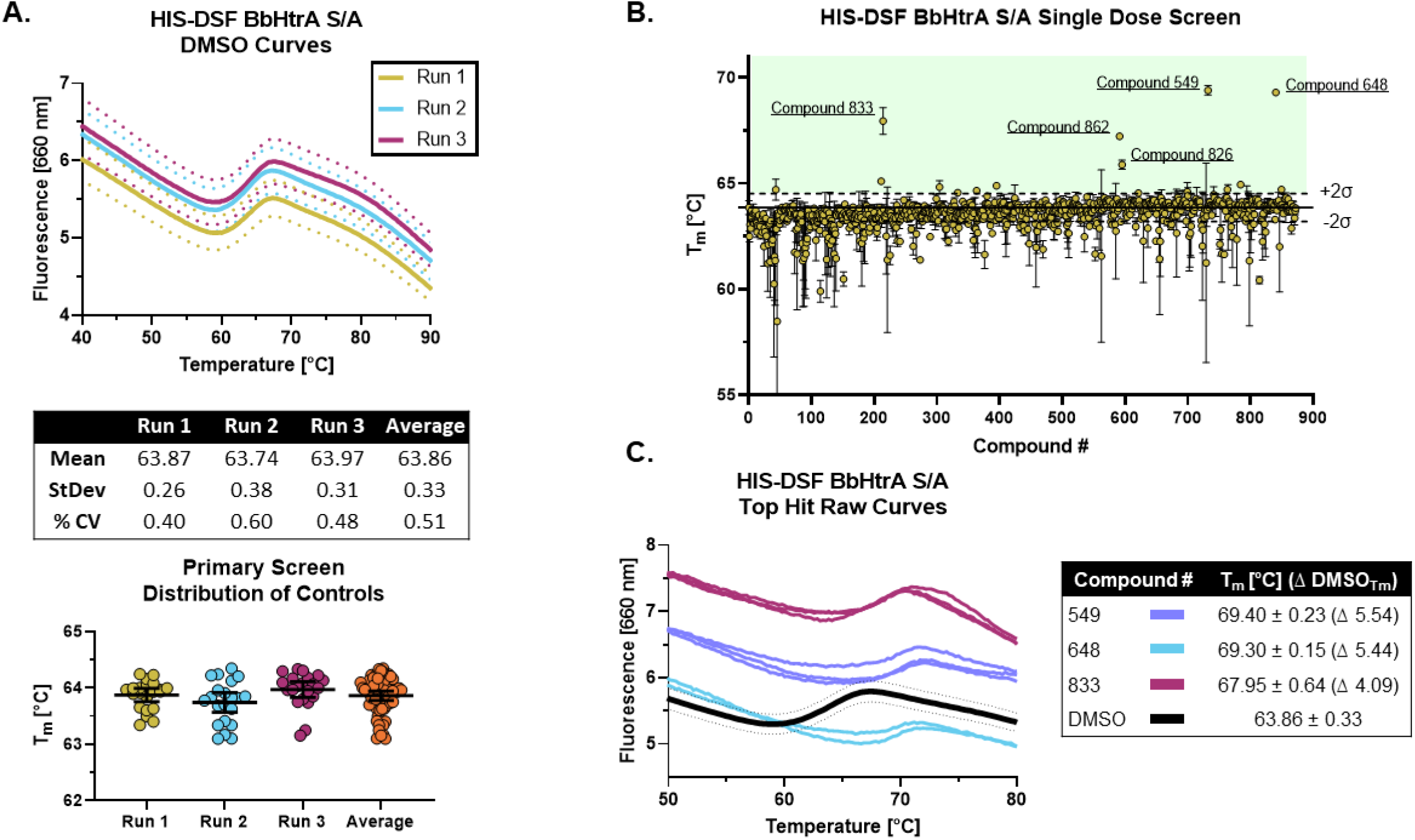
Screening of the NCATS Protease Inhibitor small molecule library using the optimized HIS-DSF assay. (A) Thermograms of the negative control DMSO wells from the three screening replicates. Each differently colored line represents the means of 24 negative control wells in a single replicate screening run with dotted lines representing the 95% confidence interval. Negative control wells are also clustered according to their T_m_ for each screening run, with whisker plots representing the mean with error bars showing the 95% confidence interval. (B) Distribution of the T_m_ values for each compound in the small molecule library arranged by masked compound ID. Each point represents the mean of the three screening runs with error bars representing the 95% confidence interval. The solid line represents the mean of the negative control DMSO wells, with dashed lines above and below representing two times the standard deviation of the negative control DMSO wells, with the green shaded area representing the area where compounds are selected for follow-up testing. (C) Thermograms of the top three hits in the HIS-DSF primary screen. Each line represents an individual replicate from each screening run, while the DMSO thermogram line represents the mean of the negative control wells with dotted lines representing the 95% confidence interval.

The cutoff for hits was made by selecting compounds that have T_m_ values greater than two times the standard deviation of the DMSO negative control groups. Using the triplicate run average T_m_ of 63.86 °C ± 0.33, compounds with an average T_m_ at or above 64.52 °C were flagged as potential hits. Applying this cutoff to the primary HIS-DSF screen filtered the 872-compound library down to 16 hits, representing a primary screen hit rate of 1.83% (Fig 4B). The thermal unfolding curves for three of these hits (**549**, **648**, and **833**) reveal a consistent right-shift in the sigmoidal unfolding curves and T_m_ of BbHtrA S/A, indicating small-molecule-ligand-induced stabilization of the target (Fig 4C). The average T_m_ for these compounds ranged from 64.54 to 69.40 °C with a median standard deviation of 0.13 °C and a % CV of 0.20%. Variation was similar when analyzing the entire single-dose library screen which had a median standard deviation of 0.17 °C and 0.26 % CV.

#### Validation and counter-screening of primary screen hits

Compounds that met the cutoff from the primary screen were replated from powder stocks and tested in a 7-point 1:4 dose response curve [200 μM – 12.8 nM final concentrations] with the same HIS-DSF assay using 5 biological replicates. The T_m_ and variation in the DMSO negative control samples was in line with previous runs (average T_m_: 63.71 °C, standard deviation: 0.28 °C, 0.44% CV) despite the higher percentage of DMSO (2% DMSO final). Of the 16 compounds that were selected from primary screening, 8 compounds demonstrate either dose-response or top-dose stabilization of BbHtrA S/A, resulting in a 50% confirmation rate from the primary screening campaign (Fig 5A). Four of the confirmed compounds had full dose-response curves with upper and lower asymptotes, allowing for robust calculation of EC_50_ values using four-parameter log-logistic fits. Three hit molecules had HIS-DSF EC_50_ values below 10 μM (**549**: 6.41 μM, **833**: 1.60 μM, and **826**: 3.64 μM) (Fig 5A), while a fourth compound, **648** had an EC_50_ value of 25.86 μM. The remaining four confirmed compounds all had stabilization at the 200 μM concentration of drug that was above the filtering criteria (T_m(DMSO)_ + 2σ_(DMSO)_) but we were unable to derive an EC_50_ value.

**Fig 5.**
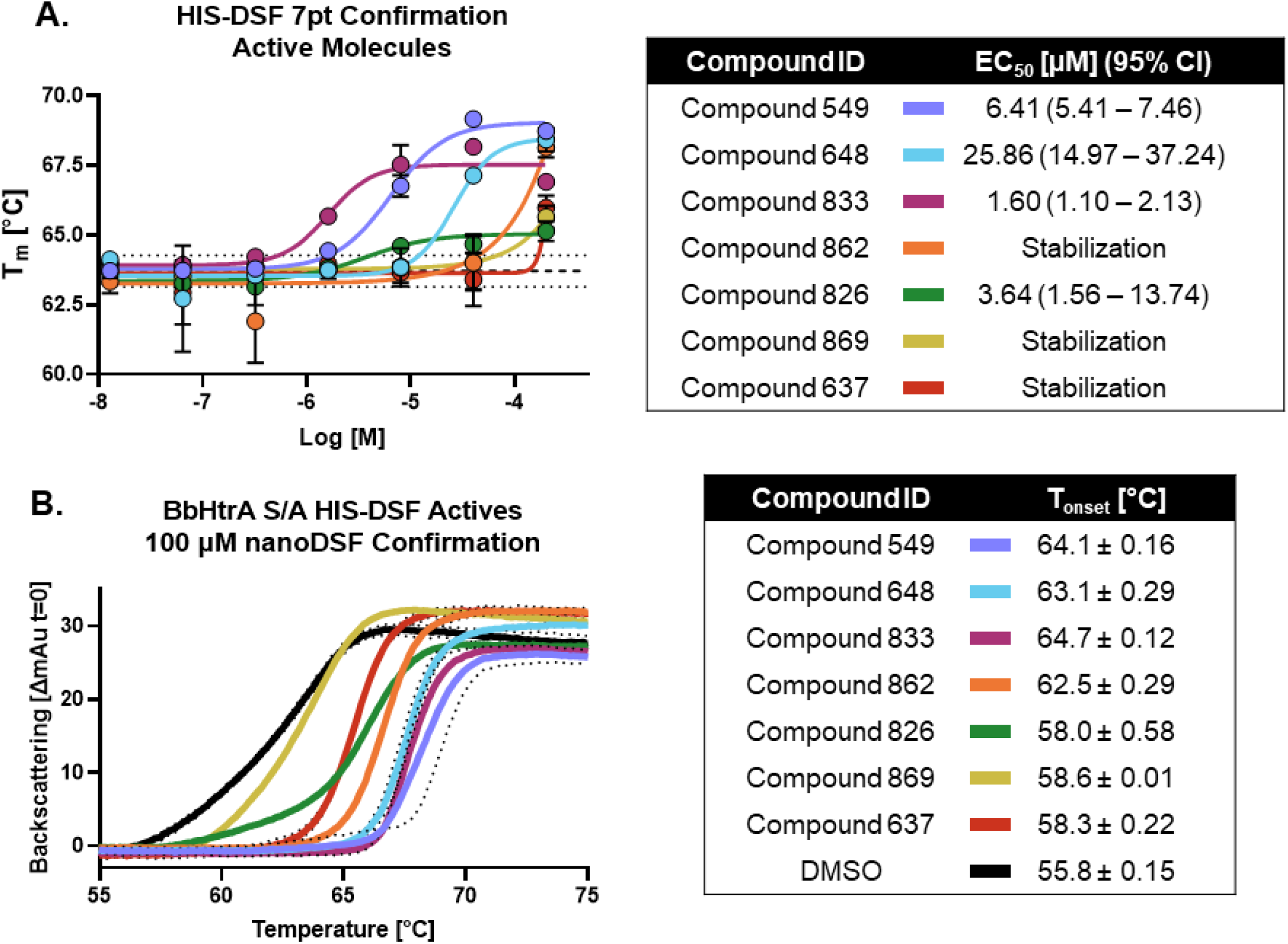
Confirmation and follow-up on the BbHtrA S/A hit molecules from the HIS-DSF primary screen. (A) Dose-response curves for the compounds that validated from the primary screen. Each point shown is the mean for five replicates at an individual concentration, with error bars representing the standard deviation of the replicates. The dose-response values were fit with a four-parameter log-logistic fit to derive the EC_50_ and 95% confidence interval. (B) Single dose stabilization as detected by nanoDSF for the 7 compounds that confirmed by dose-response HIS-DSF. The lines represent the mean backscattering signals for 3 replicates, with dotted lines representing the standard deviation of the mean.

**Fig 6.**
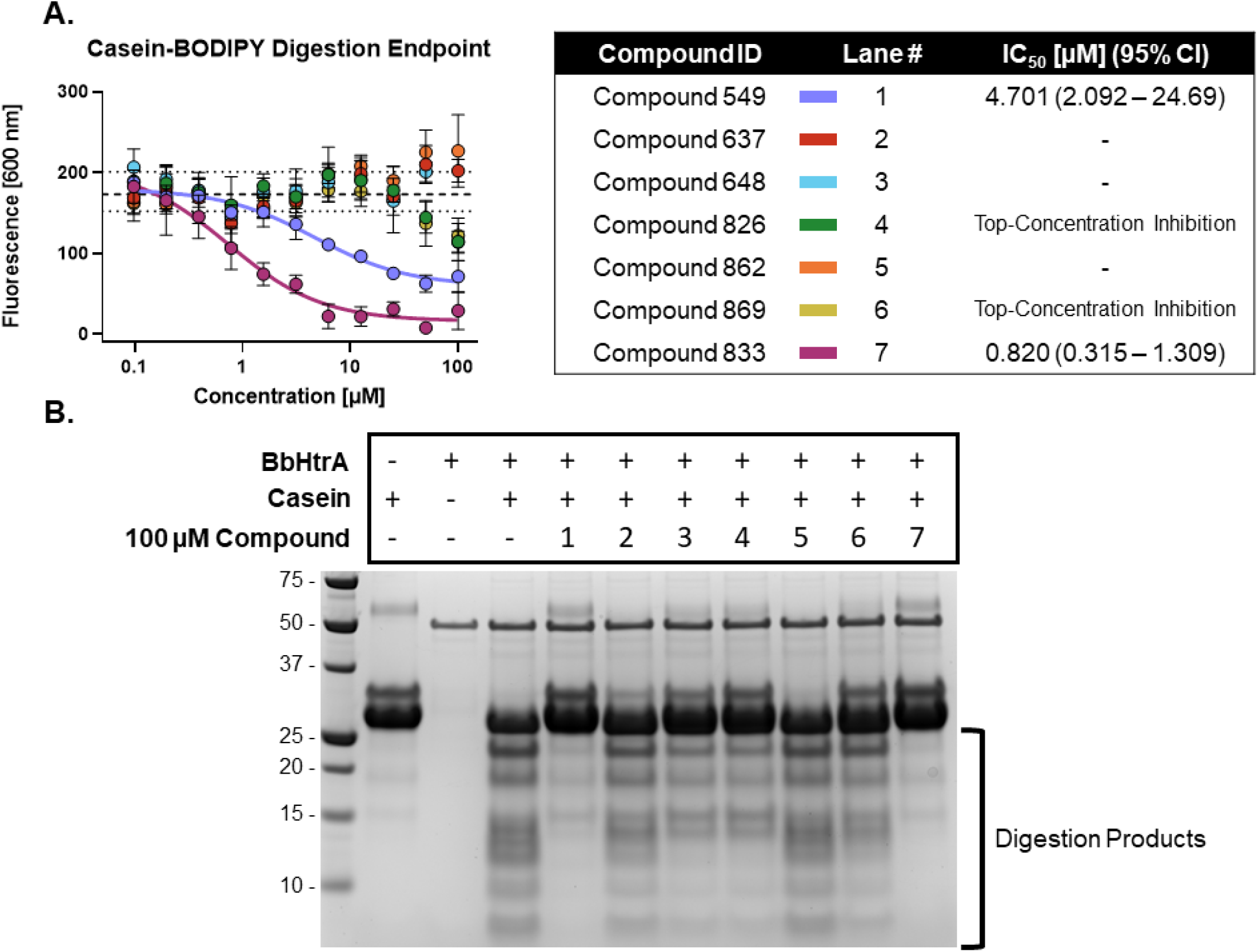
BbHtrA proteolytic activity in the presence of hit molecules. (A) Dose-response curves for hit molecules in a casein-BODIPY proteolysis assay using BbHtrA WT. Points represent the mean at each individual dose with error bars representing the standard deviation, with a dashed line and dotted lines representing the mean and ±2 standard deviations of negative control samples. The dose-response values are fit with a four-parameter log logistic fit to derive the IC_50_. (B) SDS-PAGE gel of casein digests with BbHtrA WT in the presence of 100 μM compound. Gels were stained with Imperial Blue protein stain overnight.

#### Confirmation of thermal shift using nanoDSF

In order to validate findings from the HIS-DSF assay, the 8 compounds that confirmed in 7-pt dose-response using HIS-DSF were tested using an orthogonal thermal shift-based method called nanoDSF, a differential scanning fluorimetry method that monitors shifts in tryptophan autofluorescence and backscattering aggregation signals during a thermal ramp [7, 8]. Of the 8 compounds tested at 100 μM, 7 demonstrated a significant increase in the T_onset_ as compared to the negative control (DMSO T_onset_ 55.8 °C ± 0.15) (Fig 5B). This represents a confirmation rate of 87.5% from the validated HIS-DSF hits, and an overall hit rate of 0.8% for the entire primary screen.

#### Testing hit molecules for proteolytic inhibition

Hit molecules were further profiled for their ability to inhibit the protease activity of BbHtrA by monitoring the proteolytic digestion of a model substrate, casein. Compounds were tested in dose-response using a casein substrate that has been labeled with a molar excess of BODIPY dye, resulting in a quenched substrate that fluoresces only after proteolytic cleavage. Notably, hit molecules **549** and **833** demonstrate inhibition of BbHtrA proteolysis with an IC_50_ of 4.70 and 0.820 μM, respectively. Compounds **826** and **869** also demonstrate partial inhibition at higher concentrations, but we were unable to fit a log logistic curve to the data.

Further confirmation of proteolytic inhibition was given by testing native casein digestion in the presence of inhibitors and separation of digestion fragments by SDS-PAGE. The lack of casein cleavage products in compound lane 1 and 7 signifies near total inhibition of casein proteolysis with hit molecules **549** and **833** (Fig 7B). **648**, **826**, and **869** all display partial inhibition of proteolysis, while molecules **637** and **862** don’t appear to inhibit proteolytic activity of BbHtrA (Fig 7B).

## 4 Discussion

Here, we have screened a library of 872 compounds that are known to target different classes of proteases for their ability to thermally stabilize BbHtrA. We applied a single-concentration approach to our primary screening and filtered hits by applying a selection criterion based on the standard deviation of the negative control that resulted in a hitlist of 16 compounds, a 1.6% hit rate in line with commonly reported HTS primary screens. The thermal-stabilizing effect of these hit compounds was confirmed in dose-response using the HIS-DSF assay, of which 50% were confirmed from the primary screen. These compounds were then confirmed independently of HIS-DSF using nanoDSF, in which 7 of the 8 compounds were shown to engage with BbHtrA. Additional characterization of these compounds in two independent caseinolytic assays shows that 5 of the 7 molecules can inhibit protease activity to varying degrees.

The DSF assay is sensitive to changes in the Gibbs free energy of the complex rather than any activity readout, and so compounds evolving from these biophysical campaigns are not necessarily bound to an active site of a target. This presents additional value for drug discovery projects and therapeutic targets that do not have any functional assay available, as well as a way of identifying new binding pockets that may develop in other therapeutic manners, such as with PROTACS. Interestingly, compounds **637** and **862** were unable to inhibit protease activity against casein, despite having confirmed engagement with BbHtrA as shown by HIS-DSF and nanoDSF. This lends further power to the assay in identifying binders that are potentially acting outside of the active site of a protein.

By conjugating an environment-sensitive fluorophore to the NTA moiety, there is a considerable expansion to the applicability of the DSF assay. The use of detergents and other buffer additives with SYPRO Orange results in a false signal when the dye is shuttled into the hydrophobic milieu of the surfactant micelles, an effect not seen with the NTA fluorophore employed in this study. This is particularly important in a high throughput setting where detergent use is practically necessary to prevent sticking to microfluidic lines and plateware. Additionally, the affinity-directed fluorophore will be considerably less sensitive to contaminating species present in solution, potentially allowing for measurements in the presence of cofactors, binding partners, or even in a cellular lysate.

## 5 Author Contributions

MR, BB, and AS designed the experiments and wrote the manuscript. MR and BB conducted experiments and analyzed and interpreted the data. MR and ZI worked to adapt instrumentation and create the workflows. MR, SJ, and AZ worked to create analysis scripts and methods to analyze raw data. UP and AS provided supervision and funding.

## 6 Funding

This research was supported by the Intramural Research Program of the NIH, National Center for Advancing Translational Sciences (NCATS) (ZIA TR000302-02 to A. Simeonov).

## 7 Conflict of Interest

The authors declare that the research was conducted in the absence of any commercial or financial relationships that could be construed as a potential conflict of interest.

## Notes

### Competing Interest Statement

The authors have declared no competing interest.

## References

1. Pantoliano, M.W., et al., High-density miniaturized thermal shift assays as a general strategy for drug discovery. J Biomol Screen, 2001. 6(6): p. 429–40.

2. Gao, K., R. Oerlemans, and M.R. Groves, Theory and applications of differential scanning fluorimetry in early-stage drug discovery. Biophys Rev, 2020. 12(1): p. 85–104.

3. Baljinnyam, B., et al., Applications of Differential Scanning Fluorometry and Related Technologies in Characterization of Protein–Ligand Interactions, in Targeting Enzymes for Pharmaceutical Development: Methods and Protocols, N.E. Labrou, Editor. 2020, Springer US: New York, NY. p. 47–68.

4. Ronzetti, M., et al., Protein Refolding Guided by High-Throughput Differential Scanning Fluorimetry: a case study of an HtrA-Family Bacterial Protease. bioRxiv, 2022: p. 2022.01.27.477556.

5. Bernegger, S., et al., A novel FRET peptide assay reveals efficient Helicobacter pylori HtrA inhibition through zinc and copper binding. Sci Rep, 2020. 10(1): p. 10563.

6. Russell, T.M., et al., The salt-sensitive structure and zinc inhibition of Borrelia burgdorferi protease BbHtrA. Mol Microbiol, 2016. 99(3): p. 586–96.

7. Magnusson, A.O., et al., nanoDSF as screening tool for enzyme libraries and biotechnology development. FEBS J, 2019. 286(1): p. 184–204.

8. Krakowiak, J., M. Krajewska, and J. Wawer, Monitoring of lysozyme thermal denaturation by volumetric measurements and nanoDSF technique in the presence of N-butylurea. J Biol Phys, 2019. 45(2): p. 161–172.

